# A whole-brain analysis of functional connectivity and immediate early gene expression revealed functional network shifts after operant learning

**DOI:** 10.1101/2023.10.26.564278

**Authors:** Kazumi Kasahara, Keigo Hikishima, Mariko Nakata, Tomokazu Tsurugizawa, Noriyuki Higo, Kenji Doya

## Abstract

Previous studies of operant learning have addressed neuronal activities and network changes in specific brain areas, such as the striatum, sensorimotor cortex, prefrontal/orbitofrontal cortices, and hippocampus. However, how changes in the whole-brain network are caused by cellular-level changes remains unclear. We combine resting-state functional magnetic resonance imaging (rsfMRI) and whole-brain immunohistochemical analysis of early growth response 1 (EGR1), a marker of neural plasticity, to elucidate the spatiotemporal functional network changes and underlying cellular processes during operant learning. We used an 11.7 Tesla scanner and whole-brain immunohistochemical analysis of EGR1 in mice during the early and late stages of operant learning. In the operant training, mice received a reward when they pressed the left and right buttons alternately and were punished with a bright light when they made a mistake. Control mice spent the same time and received the same amount of reward in the same operant box. A group of mice (n = 22) underwent the first rsfMRI before behavioral sessions, the second after 3 days of sessions (early stage), and the third after 21 days of sessions (late stage). Another group of mice (n = 40) was subjected to histological analysis 15 min after the early or late stages of behavioral sessions. After the early stage of training, functional connectivity was increased between the limbic areas and thalamus or auditory cortex, and the correlations of the number of EGR1-immunopositive cells between the limbic area and auditory cortex were also increased. After the late stage of training, the increases in functional connectivity and correlations of EGR1-immunopositive cells primarily occurred between the motor cortex, somatosensory cortex, and striatum. The subcortical networks centered around the limbic areas that emerged in the early stage have been implicated in rewards, pleasures, and fears. The connectivity between the motor cortex, somatosensory cortex, and striatum that consolidated in the late stage have been implicated in motor learning. Our multimodal longitudinal study successfully revealed the temporal shifts of brain regions involved in behavioral learning together with the underlying cellular-level plasticity between these regions for the first time. Our study represents a first step toward establishing a new experimental paradigm that combines rsfMRI and immunohistochemistry for linking macroscopic and microscopic mechanisms of learning.

## Introduction

When humans and animals learn a new skill, their behavior is optimized in a goal-directed manner through changes in their brains (Kolb and Whishaw, 1998). Operant conditioning, the process of acquiring new skills, is used for training animals to learn certain behaviors by associating new actions, such as lever pulling, button pressing, and nose pokes, with rewards. This has been associated with subcortical and cortical brain regions, such as the amygdala (involved in fear learning), lateral septum (involved in social playing), basal ganglia, and frontal cortex (involved in reward prediction and decision making). Enhanced activity and functional interactions in these specific regions have been demonstrated through electrophysiological experiments (Hamel et al., 2022; Zimmermann et al., 2017). However, in most animal studies, neural changes were studied in individual regions in which the electrodes were implanted. In humans, changes in the whole-brain circuits, not specific regions of interest, are frequently studied through functional magnetic resonance imaging (fMRI) (Dayan and Cohen, 2011). On the other hand, in animal studies, which allow electrophysiology and other experiments to address cellular level changes, whole-brain imaging have been used less frequently. Exceptions include a report that calculated brain-wide functional interactions via calcium imaging in zebrafish (Lin et al., 2020) and a whole-brain network map of activity-regulated gene c-Fos-positive cells in mice (Vetere et al., 2017). However, it is still unclear how whole-brain functional networks change by what cellular mechanisms in different stages of operant learning, as most techniques are invasive and cannot be observed longitudinally.

Resting-state fMRI (rsfMRI) is a non-invasive technique for evaluating brain networks with resting-state functional connectivity (FC), which can be utilized in both humans and animals for both clinical and basic research purposes. Recent studies have reported its potential utility to detect not only differences in brain connectivity between normal and diseased animal models (Tsurugizawa et al., 2020; Tudela et al., 2019; Vinh To et al., 2022), but also changes through learning tasks, leading to enhanced or inhibited task-related inter-regional connections (Sampaio-Baptista et al., 2015). Immunohistochemistry is a technique that uses antibodies to detect antigens in cells, enabling direct observation of labeled biological substances at a cellular level. However, it is not suitable for studying brain plasticity or learning in living human subjects. Early growth response 1 (*Egr1*) is a representative immediate early gene induced by neural activity during learning and plays a crucial role in long-term potentiation and synaptic plasticity (Jones et al., 2001; Veyrac et al., 2014). We hypothesized that under the same learning conditions, both immunohistochemical analysis for EGR1, the protein product of *Egr1*, and resting-state FC could be used longitudinally to reveal when, where, and how learning changes connectivity at the cellular level (i.e., EGR1 expression) and macro level (i.e., resting-state FC) during learning progresses. Here, using histological analysis of an immediate early gene and rsfMRI, we investigated spatiotemporal functional network shifts in resting-state FC and neural activity before and after short-term (early stage) and long-term (late stage) training.

## Materials and Methods

### Experimental animals

Adult male C57/BL6N mice (n = 64; 7–18 weeks old; main training or no-training started at 9 weeks old; CLEA Japan Inc., Tokyo, Japan) were used in this study. Mice were housed individually in a temperature-controlled room with 12 h light/12 h dark cycles. All behavioral training and imaging experiments were conducted during the dark cycle. All interventions and animal care procedures were conducted in accordance with the relevant guidelines and regulations and were approved by the Ethics Committee of the Okinawa Institute of Science and Technology Graduate University (approval number: 2017-178) and the National Institute of Advanced Industrial Science and Technology (approval number: 2020-0362). All animal experimental procedures were performed in accordance with the Laboratory Animal Welfare Act and the Guide for the Care and Use of Laboratory Animals (National Institutes of Health, Bethesda, MD, USA).

### Behavioral training

Six groups of mice were used to observe temporal and spatial changes in the whole-brain FC and cellular-level responses with operant conditioning. Two groups were subjected to MRI (n = 13 for training and n = 9 for no-training), and four groups were subjected to immunohistochemical analyses of ERG1 (for 21 days of training: n = 10 (training) and n = 10 (no-training); for 3 days of training: n = 10 (training) and n = 10 (no-training)) (**Fig. 1a**). Acclimation was started 3 days after water restriction at 1 mL/day (Komiyama et al., 2010). During water restriction, mice had free access to food. Water restriction and behavioral training were performed on weekdays, and water restriction was eased (to 3–5 mL/day) on weekends during which training was not performed. Mouse weight was checked daily to avoid dehydration, and water restriction was eased in case of weight loss. The MRI group underwent the first MRI before behavioral training or no-training, the second MRI after 3 days of behavioral training, and the third MRI after another 18 days of behavioral sessions. The histology group was perfused either after 3 or 21 days of behavioral sessions without MRI. Behavioral session was performed every day until day 3, every other day from days 4 to 13, and every 2 days from days 14 to 21. It took an average of 49 (range: 43–55) days to complete the training, except for the early histology group.

**Figure 1.**
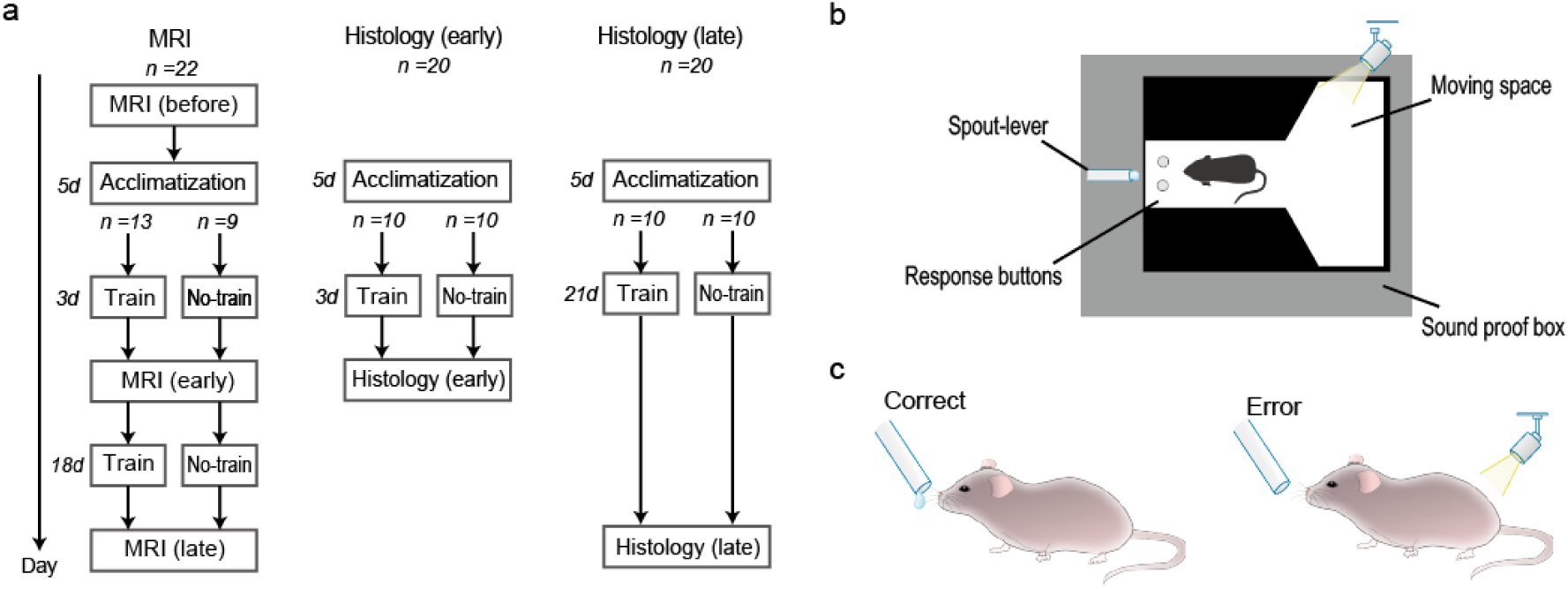
A schematic diagram of the experimental design and the behavioral task. (a) All 62 mice were acclimated to the operant unit for 5 days before the behavioral sessions. MRI was performed three times (before and after 3 days and 21 days of behavioral sessions) on 22 mice, of which 13 were subjected to behavioral training for 21 days (Train), and the remaining nine were placed in the operant unit for the same duration without training as control (No-train). Immunohistochemistry was performed on 40 mice, which were subjected to behavioral training for 21 days (n = 10) and 3 days (n = 10), or no training for 21 days (n = 10) and 3 days (n = 10). (b) The operant unit had two buttons, a waterspout, and a bright light, and was installed in a soundproof box. (c) For a correct response of pressing left and then right button, mice were rewarded with 4 µL of sugar water per trial. For an incorrect response, mice were punished with a flash of light and waiting time of 2 s.

Behavioral sessions were performed using a system installed in a soundproof box (**Fig. 1b**). Mice that responded correctly by first pressing the left button and then the right button were given sucrose water, whereas those that responded incorrectly were punished with a flash of light and a waiting period of 2 s (**Fig. 1c**). After a correct response, the next trial was immediately started. After an incorrect response, the same trial was repeated until the correct response was achieved. A training session was completed after 100 correct responses or terminated after a maximum of 1 h even if 100 correct responses were not achieved. The no-training group stayed in the operant unit for the same duration as the training group and received the same amount of sucrose water as the training group, but without training.

The mean duration and number of trials until the completion of 100 correct responses were calculated for each training day and were subjected to a one-way repeated measures analysis of variance (ANOVA) using SPSS v28.0 (SPSS, USA). Post-hoc tests with paired t-tests were performed where appropriate.

### MRI

#### Data acquisition

Data from rsfMRI were obtained using a Bruker BioSpec 117/11 11.7 Tesla MRI scanner (Bruker BioSpin GmbH, Ettlingen, Germany) with a cryogenic quadrature radio frequency surface probe (CryoProbe; Bruker BioSpin AG, Fällanden, Switzerland) to improve sensitivity. The respiration and rectal temperature of the animals were monitored during measurements. A B0 shimming method was used to improve image quality with field mapping (MAPSHIM, ParaVision 6.0.1). Data from rsfMRI were acquired with the following parameters: gradient echo-echo planer imaging sequence; repetition time/echo time = 2000/15 ms, segments = 1, FOV = 16 × 16 mm^2^, matrix = 80 × 80, slice thickness = 0.4 mm, slice gap = 0.1 mm, slice number = 20, flip angle = 50°, repetition number = 300, and scan time = 10 min. All mice were initially anesthetized with 4% isoflurane, with 2% isoflurane during set-up on the animal bed, and 1.2% isoflurane with air for rsfMRI.

#### Data analysis

The rsfMRI data analysis was performed using SPM12 (Wellcome Trust Centre for Neuroimaging, UCL Institute of Neurology, London, UK) and tailored software in MATLAB (Mathworks, Natick, MA, USA), which included adjustments for the timing of slice acquisition, motion correction, and co-registration of different mouse brains to a stereotaxic brain template (Hikishima et al., 2017). The normalized functional images were smoothed with a 0.6 mm full width at half maximum filter. A FC analysis was performed using CONN. A temporal band-pass filter was applied in the 0.009–0.1 Hz range. Excluding the cerebellum, 142 regions were used, based on an atlas (Tsurugizawa and Yoshimaru, 2021) from the Allen Institute. A functional correlation was calculated based on temporal correlation of the blood oxygenation level-dependent signal by region of interest (ROI) (**Figure 2a**).

**Figure 2.**
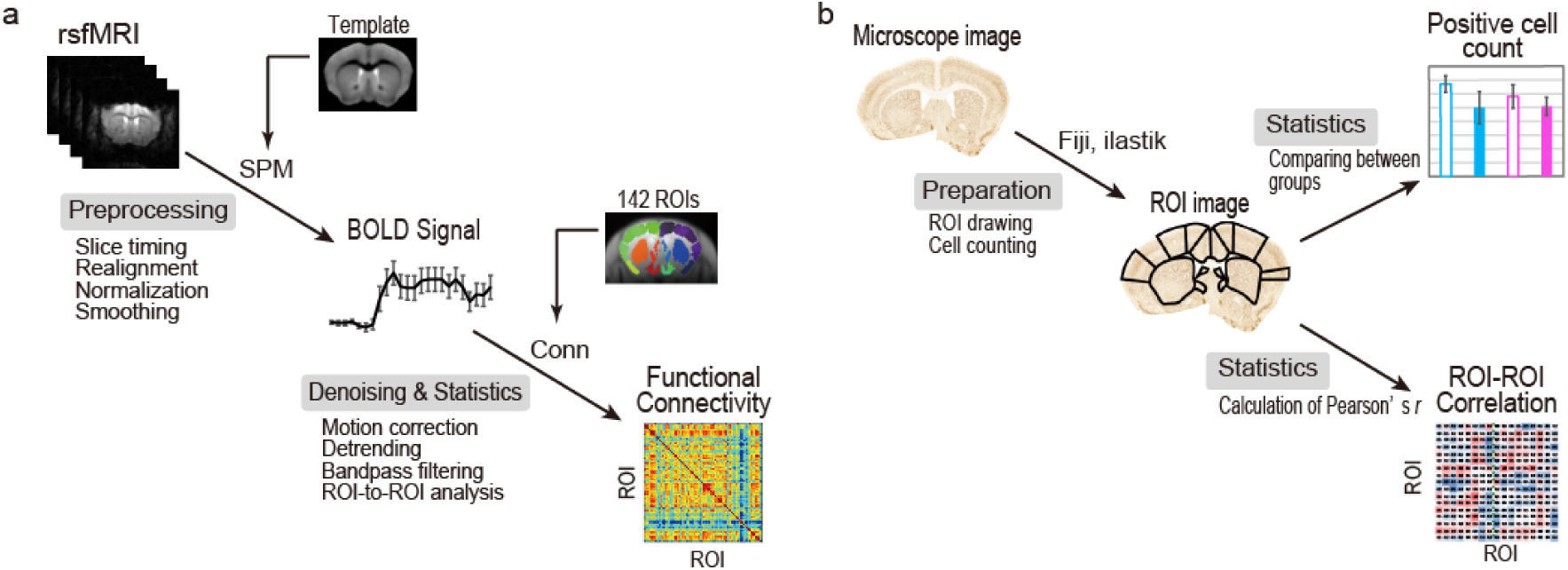
A schematic diagram outlining how to calculate resting-state functional connectivity and inter-regional correlations in immunopositive cell counts. (a) Resting-state functional data were preprocessed with SPM12, and correlations between the regions of interest (ROIs) were calculated with CONN. (b) Microscopic images were segmented into ROIs via ilastik, and positive cells were counted with Fiji. Positive cell counts were compared between groups, and correlations of the counts between ROIs were calculated. The lower right side of the brain was cut out as a “Right” marker. Abbreviations: rsfMRI, resting-state functional MRI; BOLD, blood oxygenation level-dependent.

The shifts in the resting-state FC from before training to after training (both after 3 days and after 21 days) were compared between the training and no-training groups. Second-level analyses in these shifts between training and no-training were conducted through the CONN toolbox and false discovery rate (FDR) corrected (*P* < 0.05, two-tailed).

### Histology

#### Preparation of brain tissues

Mice were deeply anesthetized with an anesthetic mixture of medetomidine (0.75 mg/kg, i.p.), midazolam (4 mg/kg, i.p.), and butorphanol (5 mg/kg, i.p.) 15 min after the end of the final trial of training or no-training (3 days for 20 mice and 21 days for 20 mice). They were then perfused with a peristaltic pump through the left cardiac ventricle with 40 mL of 100 mM phosphate buffered saline (PBS), pH 7.2, for blood removal, followed by 40 mL of 4% paraformaldehyde in PBS, pH 7.2, for fixation. Brains were removed and postfixed in the same fixative at 4 °C for 24 h. After cryoprotection in 30% sucrose in 100 mM PBS at 4 °C, coronal sections (30 μm thickness) were prepared using a cryostat. Serial sections were collected in sets of four at 120 μm intervals and stored in an anti-freezing buffer (30% ethylene glycol and 30% glycerol in 0.01 M PBS, pH 7.2, at −20 °C until use).

#### Immunohistochemistry

Sections were immunostained for EGR1. Freely floating sections were incubated in PBS containing 0.2% triton X (PBS-X) with 0.3% H_2_O_2_ for 20 min at room temperature (RT) for blocking. After washing, sections were pretreated with Block Ace (a blocking agent made from skim milk; Yukijirushi Nyugyo/Sumitomo Dainippon Pharma, Tokyo, Japan) in PBS-X (blocking buffer) for 2 h at RT. The sections were then incubated with rabbit monoclonal anti-EGR antiserum (1:2000; 15F7, Cell Signaling) in a blocking buffer overnight at 4 °C. They were washed and incubated with biotinylated goat anti-rabbit secondary antiserum (1:250; BA-1000, Vector Laboratories) in a blocking buffer for 2 h at RT. After washing, sections were incubated with the avidin-biotin complex (Vectastain ABC Elite kit, Vector Laboratories) in PBS for 1 h at RT and then washed. They were then incubated in 0.02% diaminobenzidine and 0.003% H_2_O_2_ in PBS for 2 min, and then washed with PBS.

Two-hundred and sixteen sections (−4.02 to 2.46 mm from bregma), excluding the olfactory bulb and cerebellum, were selected for histological analysis of EGR1-immunopositive cells. Each brain region that had significant resting-state FC (primary motor cortex (M1), secondary motor cortex (M2), striatum (Str), primary somatosensory upper limb (S1-ul), lateral septum ventral (LSv), lateral septum dorsal (LSd), central nucleus of the amygdala (CEA), and dorsal auditory cortex (AUDd)) was captured with a digital camera mounted to a microscope (BZ-X810, KEYENCE, USA) (**Figure 2b**). In these sections, we counted the number of EGR1-immunopositive cells in each side of the hemisphere within the targeted region with ilastik (v.1.4.0rc8, https://www.ilastik.org/) and ImageJ Fiji (v.2.9.0, https://fiji.sc). The data were analyzed in each region separately by one-way ANOVA and Bonferroni post-hoc tests between training and no-training groups. Statistically significant differences were considered at *P* < 0.05. We next computed a set of inter-regional correlations, Pearson’s *r*, across individuals in each group (Wheeler et al., 2013). The statistic t was then computed to reveal significant differences in the correlation between training and no-training in the early stage and late stage (|t | > 2.306, *P* < 0.05, two-tailed). This enabled us to identify brain regions in which EGR1 expression co-varied depending on training.

## Results

### Behavioral training

Both the duration (F_(20,380)_ = 2.964, *P* = 0.00003) and number of trials (F_(20,380)_ = 2.684, *P* = 0.00014) to achieve 100 correct responses were significantly different across the training days (**Fig. 3a**). Post-hoc analysis revealed significantly shorter duration (early: *P* = 0.004, *t* = 3.069; late: *P* = 0.003, *t* = 3.291) and fewer trials (early: *P* = 0.0005, *t* = 3.869; late: *P* = 0.0003, *t* = 4.295) in both the early stage (day 3) and late stage (day 21) compared to day 1.

**Figure 3.**
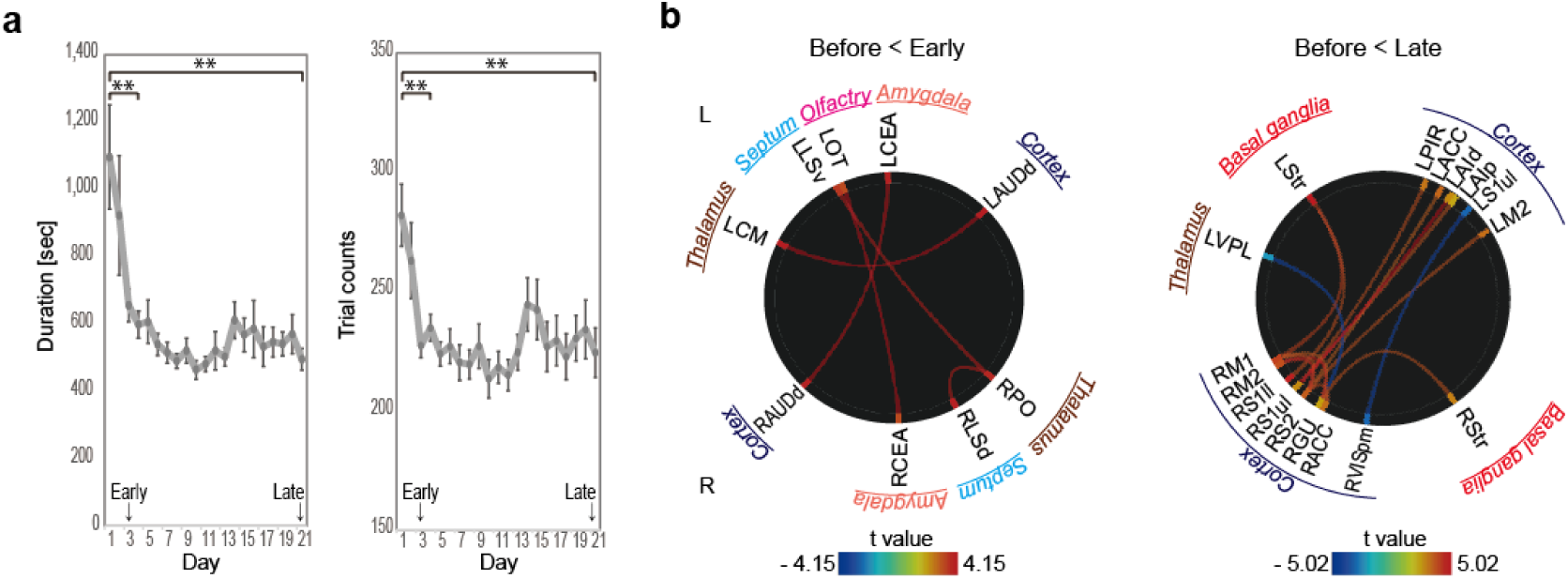
Shifts in behaviors and resting-state functional connectivity. (a) Changes in session duration and trial number to achieve 100 correct responses were significant (\*\**P* < 0.001 by one-way ANOVA or post-hoc paired t-test). (b) Short-term training (3 days; early) enhanced resting-state functional connectivity in the auditory cortex and subcortical regions (thalamus, amygdala, septum, and olfactory), and long-term training (21 days; late) enhanced resting-state functional connectivity in the cortex (motor, somatosensory, anterior cingulate, and gustatory) and striatum. Long-term training, by contrast, decreased resting-state functional connectivity between the thalamus and gustatory cortex and visual and somatosensory cortex. Central medial nucleus of the thalamus (CM), posterior complex of the thalamus (PO), lateral septal nucleus ventral (LSv), lateral septal nucleus dorsal (LSd), olfactory tubercle (OT), central nucleus of the amygdala (CEA), and dorsal auditory cortex (AUDd), ventral posterolateral nucleus of the thalamus (VPL), striatum (Str), piriform (PIR), anterior cingulate (ACC), dorsal agranular insula (AId), posterior agranular insula (AIp), primary somatosensory upper limb (S1ul), primary somatosensory lower limb (S1ll), secondary somatosensory (S2), primary motor (M1), secondary motor (M2), posteromedial visual (VISpm), gustatory (GU), L (left), and R (right).

### MRI

#### Functional connectivity

Using rsfMRI to examine changes in whole-brain functional networks, we observed enhanced resting-state FC between the auditory cortex and subcortical regions (thalamus and amygdala) and between thalamus and septum during the early stage of training (FDR-corrected, *P* < 0.05, two-tailed) (**Table 1, Early > Before, Training > No-training; Fig. 3b, left**).

**Table 1.** Enhanced significant resting-state functional connectivity during the early and late stages of training.

| Regions | to | Regions | t-value | p-FDR |
| --- | --- | --- | --- | --- |
| **Early > Before, Training > No-training** |  |  |  |  |
| <u>positive</u> |  |  |  |  |
| L dorsal auditory cortex | - | L central medial nucleus of the thalamus | 4.15 | 0.023 |
| L olfactory tubercle | - | R central nucleus of the amygdala | 3.98 | 0.038 |
| L central nucleus of the amygdala | - | R dorsal auditory cortex | 3.91 | 0.047 |
| R posterior complex of the thalamus | - | R lateral septum ventral | 3.77 | 0.039 |
| R posterior complex of the thalamus | - | L lateral septum dorsal | 3.74 | 0.039 |
| **Late > Before, Training > No-training** |  |  |  |  |
| <u>positive</u> |  |  |  |  |
| L dorsal agranular insular cortex | - | R primary somatosensory cortex upper limb | 5.02 | 0.002 |
| R secondary motor cortex | - | R gustatory cortex | 4.47 | 0.005 |
| R secondary motor cortex | - | R secondary somatosensory cortex | 4.41 | 0.005 |
| L striatum | - | R primary motor cortex | 3.99 | 0.037 |
| R secondary somatosensory cortex | - | R anterior cingulate cortex | 3.83 | 0.025 |
| L striatum | - | R primary somatosensory cortex lower limb | 3.74 | 0.039 |
| R primary somatosensory cortex lower limb | - | L piriform cortex | 3.72 | 0.036 |
| R primary motor cortex | - | R secondary somatosensory cortex | 3.67 | 0.047 |
| R secondary somatosensory cortex | - | L secondary motor cortex | 3.66 | 0.025 |
| R gustatory cortex | - | R striatum | 3.64 | 0.026 |
| R primary somatosensory cortex upper limb | - | L posterior agranular insular cortex | 3.63 | 0.036 |
| R gustatory cortex | - | R primary motor cortex | 3.51 | 0.030 |
| R secondary somatosensory cortex | - | L anterior cingulate cortex | 3.46 | 0.031 |
| <u>negative</u> |  |  |  |  |
| R posteromedial visual cortex | - | L primary somatosensory cortex upper limb | -3.92 | 0.046 |
| R gustatory cortex | - | L ventral posterolateral nucleus of the thalamus | -3.84 | 0.026 |

R, right; L, left.

During the late stage of training, enhanced resting-state FC was observed between the cortical areas (motor, somatosensory, insular, gustatory, and cingulate) and between the cortical areas (motor, somatosensory, and gustatory) and the striatum (**Table 1, Late > Before, Training > No-training; Fig. 3b, right**). Decreased resting-state FC was observed between the posteromedial visual area and S1 upper limb and gustatory cortex and left ventral posterolateral nucleus of the thalamus during the late stage, a result not observed during the early stage of training.

#### Immunohistochemistry

Positive immunoreactive staining for EGR-1 was found in the cell nucleus (**Fig. 4a**). In the early stage, positive cells were more prevalent in layers 5/6 of the auditory cortex in the training group compared to no-training group (**Fig. 4b**). In the late stage, positive cells were found to be more prevalent in layers 2/3 and 5/6 of the primary somatosensory cortex in the training group, similarly to that during the early stage (**Fig. 4c**).

**Figure 4.**
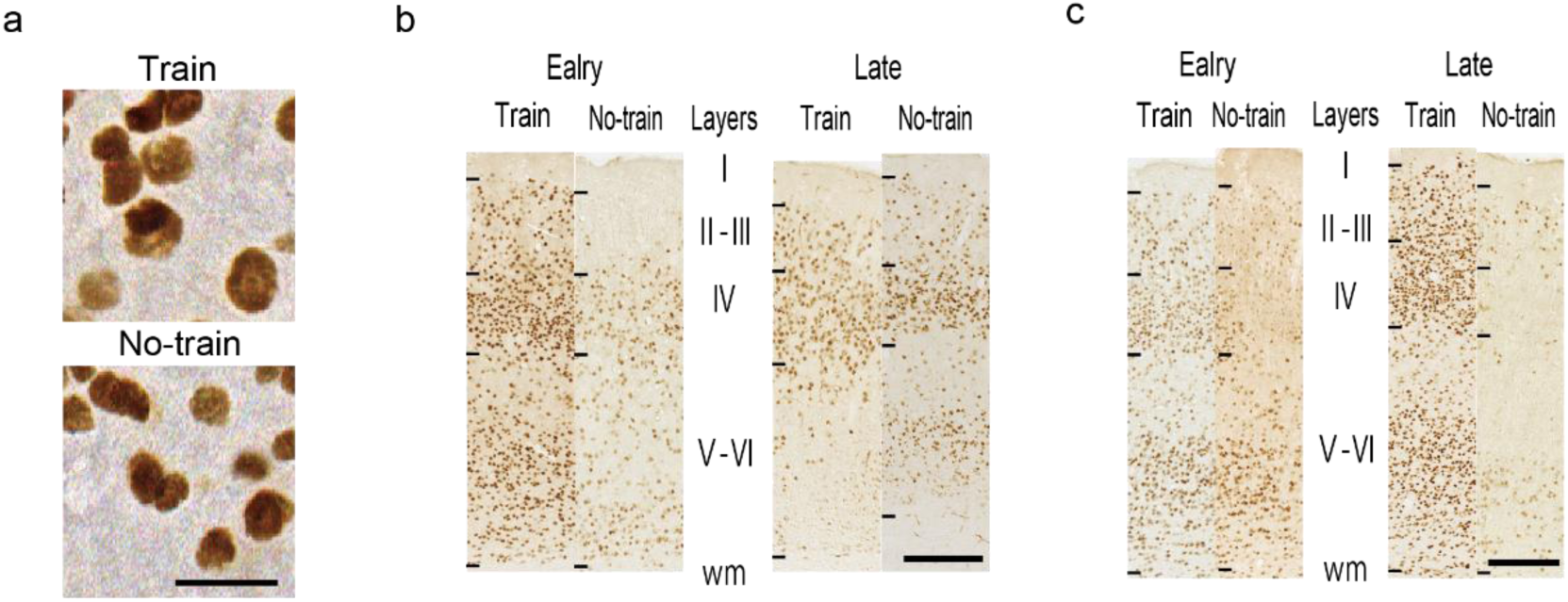
Distribution of EGR1-immunopositive cells in cortical layers. (a) Representative photomicrographs for EGR1-immunopositive cells. Both samples are from the AUDd of the late stage. Scale bar, 20 µm. Representative photomicrographs for layer distribution in (b) AUDd and (c) S1ul. During early training, positive cells were found to be more prevalent in layers 5/6 of the auditory cortex, compared to no-training and late training. During late training, positive cells were found to be more prevalent in layers 2/3 and 5/6 of the primary somatosensory cortex; however, their distribution was similar to that during early training. Scale bar, 200 µm. Dorsal auditory cortex (AUDd), primary somatosensory upper limb (S1-ul), White matter (wm).

The regions that showed changes in resting-state FC by training (Table 1) were further tested for inter-regional correlation in the numbers of EGR-1-positive cells (**Supplemental figures 1–3**). During the early stage, Pearson’s r in the training group was significantly higher than that in the no-training group between the auditory cortex and the amygdala (LAUDd-LCEA; *p* = 0.035), between motor cortices (RM1-LM1; *p* = 0.034), and between motor and primary somatosensory cortices (RM1-LS1ul; *p* = 0.022). Conversely, r between the auditory and primary somatosensory cortices (LAUDd-RS1ul; *p* = 0.076) and the auditory cortex and the striatum was significantly lower in the training group (LAUDd-RStr; *p* = 0.040) (**Fig. 5a**; not multiple-comparison-corrected).

**Figure 5.**
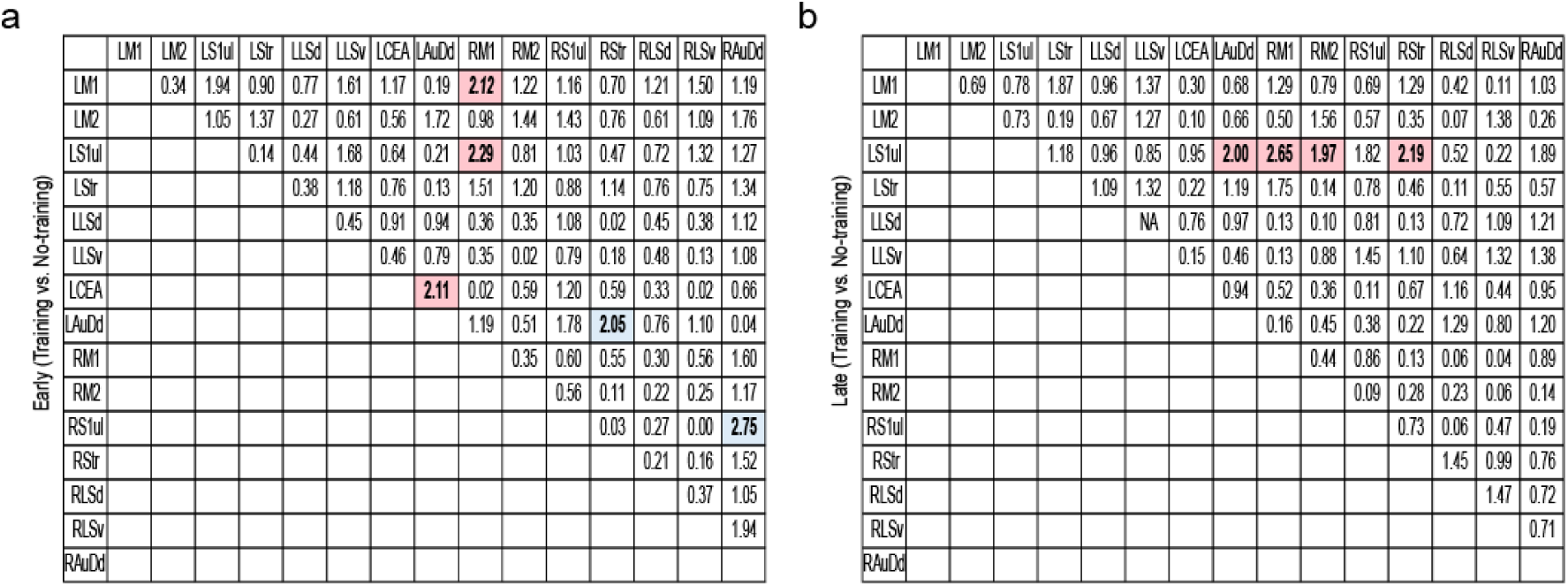
Shifts of inter-regional correlations in EGR1-positive cells. (a) Statistical z-map of the differences between no-training and training after 3 days. Significant differences in correlations of LAUDd-LCEA, RM1-LM1, RM1-LS1ul, RAUDd-RS1ul, and LAUDd-RStr between no-training and training after 3 days (*P* < 0.05, not multiple-comparison-corrected). (b) Statistical t-map of the differences between no-training and training after 21 days. Significant differences in correlations of LS1ul-RM1, LS1ul-RStr, LS1ul-RM2, and LS1ul-LAUDd between no-training and training after 21 days (*P* < 0.05, not multiple-comparison-corrected). Significant differences between correlations are highlighted in red (training has more Pearson’s *r* than no-training) and blue (training has less Pearson’s *r* than no-training) in the matrix. Dorsal auditory cortex (AUDd), central nucleus of the amygdala (CEA), primary somatosensory upper limb (S1-ul), primary motor (M1), striatum (Str), secondary motor cortex (M2), lateral septum ventral (LSv), and lateral septum dorsal (LSd).

During the late stage, Pearson’s r in the training group was significantly higher than that in the no-training group between the primary somatosensory cortex and the striatum (LS1ul-RStr; *p* = 0.029), the primary somatosensory cortex and the motor cortex (LS1ul-RM1; *p* = 0.008, LS1ul-RM2; *p* = 0.048), and the primary somatosensory cortex and the auditory cortex (LS1ul-LAUDd; *p* = 0.046). There were no regions for which Pearson’s r in the training group was lower than that for the no-training group (**Fig. 5b**; not multiple-comparison-corrected).

One-way ANOVA and post-hoc test revealed no significant differences in the number of EGR-1 positive cells in any of the regions between the four groups.

## Discussion

The present study demonstrates that multimodal investigation resting-state fMRI and immunohistochemistry can reveal the cellular-level plasticity behind changes in whole-brain dynamics. In addition, longitudinal observations before, early, and late training stages succeeded in revealing a spatiotemporal shift of network dynamics, in which subcortical networks of the limbic system related to reward were dominant in the early phase of training, whereas cortical networks related to consolidation and reconsolidation formed stronger connections in the late phase of training.

### Behavioral changes

Trial counts until mice completed the task (i.e., 100 successful trials) were nearly 300 on day 1 but decreased to approximately 220 after day 3. This indicates that mice were incorrect twice to get one correct response at the beginning of the training and only incorrect once to get one correct response after learning. This result could be interpreted in two ways. After the mice responded correctly and were rewarded once, they chose to repeat the same behavior. Conversely, the mice may have learned the rule that the opposite side was the correct side when they responded incorrectly and were not rewarded. The learning effect was observed up to day 3 and was sustained thereafter. The trial counts were slightly increased on day 14. This may be because behavioral training was performed every day until day 3, every other day from days 4 to 13, and every 2 days from days 14 to 21. Both changes in duration and trial count could include adaptation to the rules as well as improved concentration and adaptation to the environment.

### Enhanced subcortical connectivity during the early stage of training

During the early stage of training (the first 3 days), FC was enhanced between limbic areas (amygdala, lateral septum, and olfactory tubercle) and the thalamus or auditory cortex. We assumed that the enhanced connections were prevalent in the limbic area due to rewards, pleasures, and fears during the early stage of training. The central nucleus of the amygdala contributes to auditory fear conditioning with sound as a cue (Lopez et al., 2015). Nader et al. had rats remember a fear memory using an auditory fear conditioning task and then recalled the fear memory by re-exposing them to auditory stimuli (Nader et al., 2001). Furthermore, Whittle et al. reported that when CEA was suppressed by optogenetics, fear memories disappeared (Whittle et al., 2021). The olfactory tubercle is involved in feeding, feeding avoidance, and their motivations. Activation of the olfactory tubercle plays crucial roles in mediating appropriate learned (Murata et al., 2015). In this study, a beep sound was played as a cue to acquire a reward (4 μL of sugar water) upon succeeding in each trial. Hence, the enhancements in the central nucleus of the amygdala-auditory cortex and central nucleus of the amygdala-olfactory tubercle were neural circuit alterations involved in conditioning by cue sound and sugar odor. The lateral septum is responsible for reward and pleasure, as (Olds and Milner, 1954), and is (Bredewold et al., 2015; Menon et al., 2018). The posterior thalamic nucleus is involved in pain (Wang et al., 2020; Zhou et al., 2022). The posterior thalamic nucleus and lateral septum are connected via the hypothalamus. Therefore, we hypothesized that the enhanced connectivity between the posterior thalamic nucleus and lateral septum was the result of the initial response to reward and play, even though the exact role of the posterior thalamic nucleus is unknown for this task.

### Enhanced connectivity in the cortical network during the late stage of training

The enhanced FC during the late stage of training (after 21 days) primarily consisted of inter-cortical connections in the motor cortex and somatosensory cortex. Connectivity between the motor cortex, somatosensory cortex, and striatum is part of the typical motor learning circuitry, and it is likely that basal ganglia-cortical and inter-cortical connections were not fixed in the early phase of training but were established (fixed) and enhanced over time until the late phase of training. As the gustatory cortex is involved in gustatory processing, and the piriform is in direct anatomical connection with the olfactory bulb and related to odor (Stettler and Axel, 2009), we regard that the FC between the motor cortex and gustatory cortex, primary somatosensory upper limb and piriform, and striatum and gustatory cortex were enhanced as corresponding responses to this operant learning. Both the anterior cingulate and anterior insula cortex are reported to be involved in cognition and emotion, especially the former in attention and the latter in pleasure (Medford and Critchley, 2010). These cortical regions were connected to the primary somatosensory-upper limb, which controls forelimb somatosensory functions, and their connections may have been relevant in the execution and retention of the present training, which engaged the forelimbs. In the present study, we repeated training 26 times, including 5 days of habituation, 3 days of pretraining, and 18 days of training. Therefore, there was little novelty regarding the device, button, and visual, somatosensory, and gustatory rewards by the late phase. The lack of these novelties reduced FC in the visual cortex, primary somatosensory upper limb, gustatory cortex, and ventral posterolateral nucleus of the thalamus.

### Biological shifts observed during the early and late stages of training

The immediate early gene *Egr1* is expressed in a neural activity-dependent manner and is involved in memory reconsolidation (Lee et al., 2004), long-term potentiation (Jones et al., 2001), and synaptic plasticity (Veyrac et al., 2014). Newly learned memories are acquired and then fixed in a time-dependent manner; however, they are unstable and fragile, and require reconsolidation to become permanent memories. Reconsolidation involves maintaining the original memory intact and stored in the brain, enhancing the original memory, and updating the memory through integration or modification with another new memory (Trent et al., 2015). In the present study, it is possible that, as the mice learned a new skill and FC was increased, long-term potentiation was induced and the memory was enhanced through EGR1 expression. Both EGR1-regional correlation and FC were consistently increased between the amygdala and auditory cortex during the early stage of training, and between the somatosensory-motor and striatum during the late stage of training. In other words, this study suggests that neural activity shifted from subcortical to cortical regions during learning something new and was observed as increased FC. However, there was a difference between MRI and histological analyses of the bilateral sensorimotor cortex.

Histological analysis revealed that bilateral somatosensory-motor areas correlated with each other during both the early and late stages. MRI revealed the correlation during the late but not early stage. Because our training comprised left-right button pressing, we consider it reasonable that bilateral somatosensory-motor correlations would be higher at any time point, and that this was due to the difference in the target measured: synchronization of BOLD oscillation during resting state for MRI and local neuronal activation during task execution for EGR1. In the present study, we focused on EGR1, which is commonly induced during learning as a first step. However, it is also possible that a signaling pathway other than EGR1 may be involved in this learning. Future comprehensive analysis including various molecules, not only EGR1, may reveal which signaling pathway is highly related to the neural activity induced by this type of learning. These results for FC and EGR1 were approximately consistent, and it could be argued that EGR1 is one of the molecular bases underlying the FC changes observed with MRI. This study is the first step toward establishing an experimental system that combines MRI and histological analysis.

### The potential of our method that combines MRI and histological analysis

Our observations of layer-specific changes of EGR1 expression in the auditory and primary somatosensory cortices provide further insights as to the FC change in rsfMRI and cortical circuit organization (**Fig. 4**): EGR1 expression was increased in layer 5/6 of auditory cortex, connected to subcortical areas such as the amygdala, during early training and in layers 2/3, connected to other cortices and layers 5/6 of somatosensory cortex, connected to the striatum, during late training. Although the present study did not perform quantitative analysis of layer structures, this is the first step in establishing an experimental system combining MRI and whole-brain immunohistochemical analysis. Therefore, this strategy may be extended to multimodal and multi-scale analyses based on the distribution of each cortical cell type and each cortical layer to gain insight into network changes during learning.

## Limitation

We assessed neural activity during learning by analyzing EGR1 expression and rsfMRI after learning. We did not assess neural activity directly or any psychophysiological interactions during learning, and it is unclear whether the enhanced FC after learning was similarly enhanced during learning. Although it is very difficult to measure brain activity in head-fixed rodents performing tasks inside a small MRI bore, understanding functional shifts during learning may be needed in the future (Han et al., 2019). Analysis of EGR1 expression revealed differences in inter-regional correlations across subjects in training and no-training groups, but not in the number of positive cells in each region between groups. In this study, mice spent the same amount of time and received the same amount or reward in the same operant system in both the no-training and training groups. In other words, in both groups, the mice experienced the same environment except for learning the conditions for reward. Therefore, it is possible that the difference in neural activities specific to training was relatively small, potentially resulting in no significant difference in the number of EGR1-positive cells. Thus, further investigation is needed.

## Conclusion

This study combined MRI and histological analysis and successfully visualized temporal shifts in brain regions involved in behavioral learning and demonstrated the plasticity in these regions for the first time together. This study represents the first step toward establishing an experimental system combining both MRI and whole-brain immunohistochemical analysis.

## Acknowledgments

This study was funded by the Japan Society for the Promotion of Science grants KAKANHI 21K19463 and KAKANHI 20H04236 (KK) and Japan Science and Technology Agency grant FORESTO JPMJFR206G (KK).

## CRediT author statement

**Kazumi Kasahara**: Conceptualization, Methodology, Investigation, Writing—original draft, Writing—review & editing, Writing—review & editing, Funding acquisition, Resources, Supervision. **Keigo Hikishima**: Methodology, Investigation, Writing—original draft, Writing—review & editing, Supervision. **Mariko Nakata**: Methodology, Investigation, Writing—original draft, Writing—review & editing. **Tomokazu Tsurugizawa**: Writing—original draft, and Writing—review & editing. **Noriyuki Higo**: Writing—original draft, Writing—review & editing. **Kenji Doya**: Conceptualization, Methodology, Writing—original draft, Writing—review & editing, Resources.

## Notes

### Competing Interest Statement

The authors have declared no competing interest.

