## Supplemental Figure1, 2, 3 for "A whole-brain analysis of functional connectivity and immediate early gene expression revealed functional network shifts after operant learning"

Including 3 Figures.

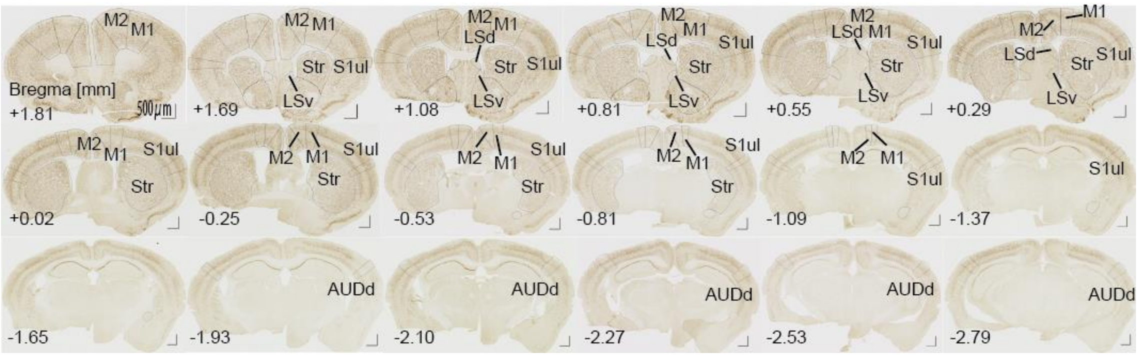


**Supplemental Figure 1. Immunohistochemistry images and regions of interest in untrained mice.**

Mice were perfused 15 min after pseudo-training (no training) on days 3 (n = 10; early stage) and 21 (n = 10; late stage). The image is an example of the late stage. Each brain was sliced every 30 µm to identify the regions of interest (ROIs) within −4.02 mm to 2.46 mm from bregma. The lower right section of the brain was removed as a marker. Abbreviations: M1, primary motor cortex; M2, secondary motor cortex; Str, striatum; S1-ul, primary sensory upper limb; LSv, lateral septum ventral; LSd, lateral septum dorsal; CEA, central nucleus of the amygdala; AUDd, dorsal auditory cortex.

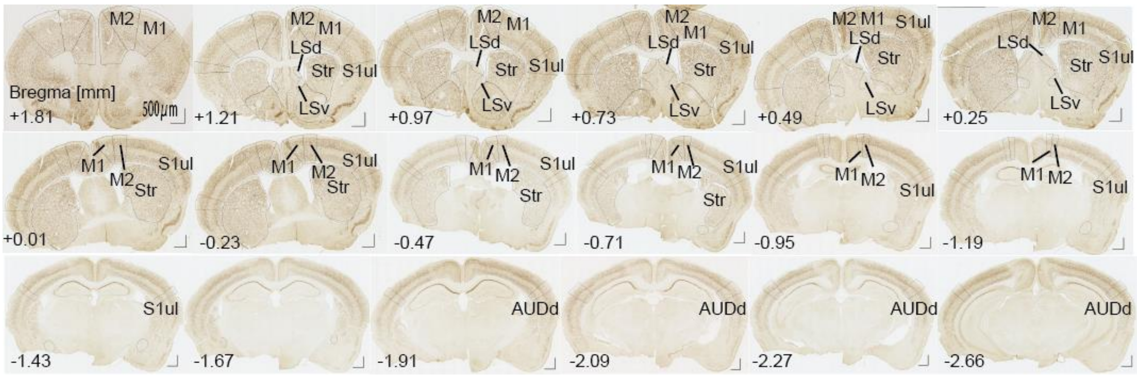


**Supplemental Figure 2. Immunohistochemistry images and regions of interest in trained mice.** Mice were perfused 15 minutes after training on the third day (n= 10; early stage) and the 21st day (n=10; late stage). This image is an example of the late stage. Each brain was sliced every 30 μm to identify the regions of interest (ROIs) within -4.02 mm to 2.46 mm from bregma. The lower right section of the brain was removed as a marker. Abbreviations: M1, primary motor cortex; M2, secondary motor cortex; Str, striatum; S1-ul, primary sensory upper limb; LSv, lateral septum ventral; LSd, lateral septum dorsal; CEA, central nucleus of the amygdala; AUDd, dorsal auditory cortex.

**a**

| | LAuDd | LM2 | LM1 | LS1ul | LStr | LLSd | LLSv | LCEA | RAuDd | RM2 | RM1 | RS1ul | RStr | RLSd | RLSv |
|---|---|---|---|---|---|---|---|---|---|---|---|---|---|---|---|
| LAuDd | | -0.47 | -0.07 | 0.08 | 0.04 | -0.20 | 0.22 | 0.48 | 0.70 | -0.13 | -0.14 | -0.32 | -0.10 | 0.02 | -0.04 |
| LM2 | | | 0.75 | 0.66 | 0.80 | 0.21 | 0.37 | -0.28 | -0.61 | 0.67 | 0.58 | 0.77 | 0.68 | 0.28 | 0.63 |
| LM1 | | | | 0.96 | 0.79 | 0.06 | 0.67 | 0.05 | -0.23 | 0.76 | 0.89 | 0.82 | 0.45 | 0.38 | 0.78 |
| LS1ul | | | | | 0.78 | 0.20 | 0.79 | 0.14 | -0.27 | 0.65 | 0.88 | 0.77 | 0.38 | 0.52 | 0.83 |
| LStr | | | | | | 0.12 | 0.47 | -0.16 | -0.19 | 0.70 | 0.50 | 0.66 | 0.72 | 0.52 | 0.67 |
| LLSd | | | | | | | 0.45 | -0.28 | -0.67 | -0.36 | 0.02 | 0.21 | 0.02 | 0.68 | 0.49 |
| LLSv | | | | | | | | -0.01 | -0.31 | 0.17 | 0.69 | 0.65 | 0.37 | 0.53 | 0.76 |
| LCEA | | | | | | | | | 0.32 | 0.27 | 0.14 | -0.32 | -0.50 | -0.23 | 0.13 |
| RAuDd | | | | | | | | | | -0.02 | -0.32 | -0.46 | -0.18 | -0.38 | -0.42 |
| RM2 | | | | | | | | | | | 0.58 | 0.47 | 0.40 | 0.07 | 0.55 |
| RM1 | | | | | | | | | | | | 0.83 | 0.26 | 0.19 | 0.61 |
| RS1ul | | | | | | | | | | | | | 0.65 | 0.25 | 0.60 |
| RStr | | | | | | | | | | | | | | 0.11 | 0.38 |
| RLSd | | | | | | | | | | | | | | | 0.61 |
| RLSv | | | | | | | | | | | | | | | |

**b**

| | LAuDd | LM2 | LM1 | LS1ul | LStr | LLSd | LLSv | LCEA | RAuDd | RM2 | RM1 | RS1ul | RStr | RLSd | RLSv |
|---|---|---|---|---|---|---|---|---|---|---|---|---|---|---|---|
| LAuDd | | 0.39 | 0.03 | -0.03 | 0.11 | 0.29 | 0.57 | -0.54 | 0.71 | 0.14 | 0.46 | 0.55 | 0.76 | 0.40 | 0.50 |
| LM2 | | | 0.66 | 0.23 | 0.35 | 0.07 | 0.06 | 0.01 | 0.23 | 0.04 | 0.14 | 0.25 | 0.40 | -0.04 | 0.16 |
| LM1 | | | | 0.72 | 0.53 | -0.34 | -0.05 | 0.59 | 0.38 | 0.33 | 0.28 | 0.49 | 0.11 | -0.24 | 0.24 |
| LS1ul | | | | | 0.75 | -0.03 | 0.17 | 0.45 | 0.38 | 0.33 | 0.15 | 0.44 | 0.15 | 0.19 | 0.45 |
| LStr | | | | | | -0.08 | -0.12 | 0.24 | 0.48 | 0.22 | -0.25 | 0.31 | 0.29 | 0.17 | 0.39 |
| LLSd | | | | | | | 0.62 | -0.65 | -0.21 | -0.51 | -0.17 | -0.35 | 0.03 | 0.79 | 0.32 |
| LLSv | | | | | | | | -0.25 | 0.25 | 0.18 | 0.58 | 0.34 | 0.45 | 0.69 | 0.73 |
| LCEA | | | | | | | | | -0.02 | 0.53 | 0.15 | 0.30 | -0.23 | -0.39 | 0.14 |
| RAuDd | | | | | | | | | | 0.54 | 0.48 | 0.75 | 0.56 | 0.16 | 0.53 |
| RM2 | | | | | | | | | | | 0.69 | 0.67 | 0.45 | -0.05 | 0.45 |
| RM1 | | | | | | | | | | | | 0.70 | 0.51 | 0.03 | 0.39 |
| RS1ul | | | | | | | | | | | | | 0.66 | 0.11 | 0.60 |
| RStr | | | | | | | | | | | | | | 0.22 | 0.45 |
| RLSd | | | | | | | | | | | | | | | 0.72 |
| RLSv | | | | | | | | | | | | | | | |

**c**

| | LAuDd | LM2 | LM1 | LS1ul | LStr | LLSd | LLSv | LCEA | RAuDd | RM2 | RM1 | RS1ul | RStr | RLSd | RLSv |
|---|---|---|---|---|---|---|---|---|---|---|---|---|---|---|---|
| LAuDd | | 0.18 | 0.32 | 0.41 | -0.09 | 0.12 | 0.22 | 0.09 | 0.83 | 0.38 | 0.40 | -0.04 | -0.03 | 0.25 | 0.14 |
| LM2 | | | 0.51 | 0.56 | 0.48 | 0.38 | 0.17 | -0.18 | -0.05 | 0.60 | 0.30 | 0.41 | 0.70 | 0.69 | 0.37 |
| LM1 | | | | 0.81 | 0.70 | 0.39 | 0.50 | -0.30 | 0.58 | 0.45 | 0.81 | 0.29 | 0.27 | 0.36 | 0.12 |
| LS1ul | | | | | 0.60 | 0.58 | 0.51 | -0.04 | 0.50 | 0.78 | 0.88 | 0.66 | 0.57 | 0.56 | 0.45 |
| LStr | | | | | | 0.74 | 0.57 | -0.26 | 0.07 | 0.24 | 0.46 | 0.42 | 0.49 | 0.56 | 0.14 |
| LLSd | | | | | | | 0.78 | -0.13 | 0.15 | 0.52 | 0.40 | 0.62 | 0.57 | 0.72 | 0.54 |
| LLSv | | | | | | | | -0.18 | 0.44 | 0.41 | 0.51 | 0.34 | 0.09 | 0.27 | 0.20 |
| LCEA | | | | | | | | | -0.14 | -0.14 | -0.28 | -0.06 | -0.26 | -0.03 | -0.15 |
| RAuDd | | | | | | | | | | 0.33 | 0.65 | -0.03 | -0.20 | 0.04 | 0.06 |
| RM2 | | | | | | | | | | | 0.73 | 0.60 | 0.64 | 0.57 | 0.66 |
| RM1 | | | | | | | | | | | | 0.42 | 0.32 | 0.25 | 0.27 |
| RS1ul | | | | | | | | | | | | | 0.80 | 0.60 | 0.78 |
| RStr | | | | | | | | | | | | | | 0.79 | 0.72 |
| RLSd | | | | | | | | | | | | | | | 0.74 |
| RLSv | | | | | | | | | | | | | | | |

**d**

| | LAuDd | LM2 | LM1 | LS1ul | LStr | LLSd | LLSv | LCEA | RAuDd | RM2 | RM1 | RS1ul | RStr | RLSd | RLSv |
|---|---|---|---|---|---|---|---|---|---|---|---|---|---|---|---|
| LAuDd | | -0.17 | -0.03 | -0.56 | -0.62 | -0.38 | -0.02 | 0.53 | 0.50 | 0.16 | 0.47 | 0.16 | 0.09 | -0.41 | -0.28 |
| LM2 | | | 0.19 | 0.24 | 0.40 | 0.64 | 0.69 | -0.13 | 0.09 | 0.91 | 0.52 | 0.63 | 0.59 | 0.71 | 0.81 |
| LM1 | | | | 0.61 | -0.13 | -0.10 | -0.18 | -0.44 | 0.11 | 0.06 | 0.41 | -0.07 | -0.39 | 0.15 | 0.06 |
| LS1ul | | | | | 0.06 | 0.15 | 0.11 | -0.50 | -0.43 | -0.01 | -0.04 | -0.18 | -0.48 | 0.34 | 0.35 |
| LStr | | | | | | 0.35 | -0.06 | -0.15 | -0.23 | 0.17 | -0.41 | 0.03 | 0.28 | 0.52 | 0.41 |
| LLSd | | | | | | | 0.78 | -0.49 | -0.46 | 0.48 | 0.34 | 0.82 | 0.52 | 0.86 | 0.83 |
| LLSv | | | | | | | | -0.10 | -0.26 | 0.72 | 0.56 | 0.81 | 0.59 | 0.55 | 0.72 |
| LCEA | | | | | | | | | 0.35 | 0.05 | -0.01 | -0.12 | 0.09 | -0.57 | -0.37 |
| RAuDd | | | | | | | | | | 0.26 | 0.29 | -0.13 | 0.20 | -0.33 | -0.31 |
| RM2 | | | | | | | | | | | 0.60 | 0.63 | 0.72 | 0.48 | 0.64 |
| RM1 | | | | | | | | | | | | 0.72 | 0.38 | 0.22 | 0.29 |
| RS1ul | | | | | | | | | | | | | 0.61 | 0.62 | 0.66 |
| RStr | | | | | | | | | | | | | | 0.29 | 0.36 |
| RLSd | | | | | | | | | | | | | | | 0.94 |
| RLSv | | | | | | | | | | | | | | | |

**Supplemental Figure 3. Significant correlations in numbers of EGR1-positive cells between regions. a**. Early stage of training. **b**. Early stage of no-training. **c**. Late stage of training. **d**. Late stage of no-training. Significant differences between correlations are highlighted in red (positive) and blue (negative). Pearson's $r$, $P < 0.05$.
